# Ribo-seQC: comprehensive analysis of cytoplasmic and organellar ribosome profiling data

**DOI:** 10.1101/601468

**Authors:** Lorenzo Calviello, Dominique Sydow, Dermot Harnett, Uwe Ohler

**Author notes:** Equal contribution (Lorenzo Calviello, Dominique Sydow).

## Abstract

**Summary:** Ribosome profiling enables genome-wide analysis of translation with unprecedented resolution. We present *Ribo-seQC*, a versatile tool for the comprehensive analysis of Ribo-seq data, providing in-depth insights on data quality and translational profiles for cytoplasmic and organelle ribosomes. *Ribo-seQC* automatically generates platform-independent HTML reports, offering a detailed and easy-to-share basis for collaborative Ribo-seq projects.

**Availability:** Ribo-seQC is available at https://github.com/ohlerlab/RiboseQC and submitted to Bioconductor.

## 1. Introduction

Ribosome profiling allows to monitor the position of translating ribosomes, providing a global and detailed view on protein synthesis (Ingolia *et al*., 2009). Analysis and visualization of such rich datasets are a challenging endeavor, and require suitable computational tools. Published methods such as *Shoelaces* (Birkeland *et al*., 2018), *riboviz* (Carja *et al*., 2017), *riboWaltz* (Lauria *et al*., 2018), *Ribo-TISH* (Zhang *et al*., 2017), *RiboProfiling* (Popa *et al*., 2016), *systemPipeR* (Backman and Girke, 2016), *RUST* (O’Connor *et al*., 2016), and *riboSeqR* (Chung *et al*., 2015) share basic functionalities, albeit differing in their primary focus, e.g. on providing general quality control or automated P-sites positions generation (see Supplementary Tab. 1). However, these methods cannot analyze footprints originating from cytoplasmic and organellar ribosomes, and often do not provide sufficiently detailed statistics on ribosome profiles on different annotated biotypes (e.g. UTRs, intergenic regions, non-coding RNAs). Moreover, summarizing the rich output of such analyses is a challenging task, often involving the users to browse multiple files with several plots for each dataset, which represents an obstacle for non-specialists.

*Ribo-seQC* aims at addressing these drawbacks. Our tool offers functionalities for quality control, including read length and biotype-specific statistics as well as abundantly mapped positions (potentially corresponding to undesired artifacts). Footprint profiles can be assessed by investigating metagene profiles and codon usage with single nucleotide resolution or bins, per each footprint length, with multiple visualization options, including profiles for organellar ribosomes. Importantly, multiple samples can be investigated in the same output, all summarized in one HTML report containing a detailed description of the output and publication-ready figures.

## 2. Implementation

*Ribo-seQC* is implemented as an R Bioconductor package, and it can (1) generate genome and annotation in R for the species under investigation, (2) analyze one or multiple Ribo-seq samples, and (3) automatically generate the comprehensive HTML report.

### 2.1 Genome and annotation generation

Genome and annotation R objects are generated from user-provided 2bit and GTF files, using *BSgenome* (Pagès, 2018) and *GenomicFeatures* (Lawrence *et al*., 2013), respectively.

### 2.2 Read analysis

BAM files are loaded and processed in small chunks, using *GenomicFiles* (Bioconductor Package Maintainer *et al*., 2018), to ensure efficient computation and limit RAM usage. Initially, all reads are split by nuclear and organellar genomes for independent processing. Reads are subsequently aligned to annotated genomic regions, using *GenomicAlignments* (Lawrence *et al*., 2013).

We report **footprint length and biotype-specific statistics**, as well as a combination of both to investigate a potential prevalence of distinct footprint lengths in certain biotypes. Moreover, we provide **count statistics per gene**, including the standard normalization measures RPKM and TPM. In order to determine contaminants in experiments, we report the **most abundant mapping positions**, together with their abundance and nucleotide sequence.

**Metagene analysis** of 5’ and P-sites profiles are calculated in two ways: (i) At sub-codon resolution, we summarize read coverage in the vicinity of start and stop codons, as well as regions in the middle of the CDS, for a detailed view of the ribosomal three-nucleotide movement. (ii) At the resolution of fixed percentile bins, we capture read coverage over coding transcripts, considering UTRs and CDS lengths separately (50 bins for 5’ and 3’UTR and 100 CDS bins). Profiles are generated for individual footprint lengths, and provided as raw, z-scored or log2-transformed counts.

Many tools set a preselected read length around 30nt and offset around 12nt to generate P-sites from 5’ profiles. However, read lengths can differ between protocols, nuclear and organellar genomes, and even biotypes. Thus, we first select all in-frame **read lengths**, i.e. read lengths showing greater coverage in one frame when compared to the off-frame signal. This is performed using average values across all transcripts, thus limiting the effect of high off-frame signal on individual transcripts. Subsequently, we automatically select **offsets** for each read length, calculating the maximum distance around start codon positions, using both aggregate and averaged profiles across transcripts. P-sites positions (calculated using frame and offset per each read length) are then used to create aggregate profiles.

We offer an extensive analysis on **codon usage**, both as bulk (transcriptome-wide) and positional (codon resolution) analysis, again distinguishing between the first, middle, and last codon positions. We provide codon counts (sequence only), P-sites counts, and P-sites-to-codon ratios.

### 2.3 Automatic report generation

*Ribo-seQC* produces a single R Markdown-based report in the platform-independent format HTML. Notably, multiple samples, nuclear and organellar genomes, as well as a wide range of read lengths can be explored interactively and comparatively. Publication-ready plots are provided.

In addition, bigwig files for read coverage and P-sites positions are automatically generated. All data used to create the entire report is made easily accessible through the use of compressed RData files.

## 3. Results

To demonstrate the utility of *Ribo-seQC*, we applied it to Ribo-seq data from *Arabidopsis thaliana* roots and shoots (Hsu *et al*., 2016). As revealed by *Ribo-seQC*, the shoot Ribo-seq datasets included extensive ribosome footprints mapping to the chloroplast genome (Fig. 1a). An example P-sites profile for 29nt chloroplast footprints in the shoot sample is shown in Fig. 1b. Footprint length-specific frame preference in the root sample (Fig. 1c) demonstrates again the importance of performing organelle-specific and also read length-specific analysis of ribosomal footprints. Positional analysis of codon usage (P-sites per codon) in the root sample is shown in Fig. 1d, separating start and stop codons from other codons. Bulk analysis of codon usage is also shown for the shoot sample in Fig. 1e. Analysis of top mapping positions (Fig. 1f) reveals the clear presence of abundant ribosomal RNA contamination; the footprint sequence, together with the gene IDs they map to and the percentage of library they represent, are also provided for the different samples.

**Fig. 1.**
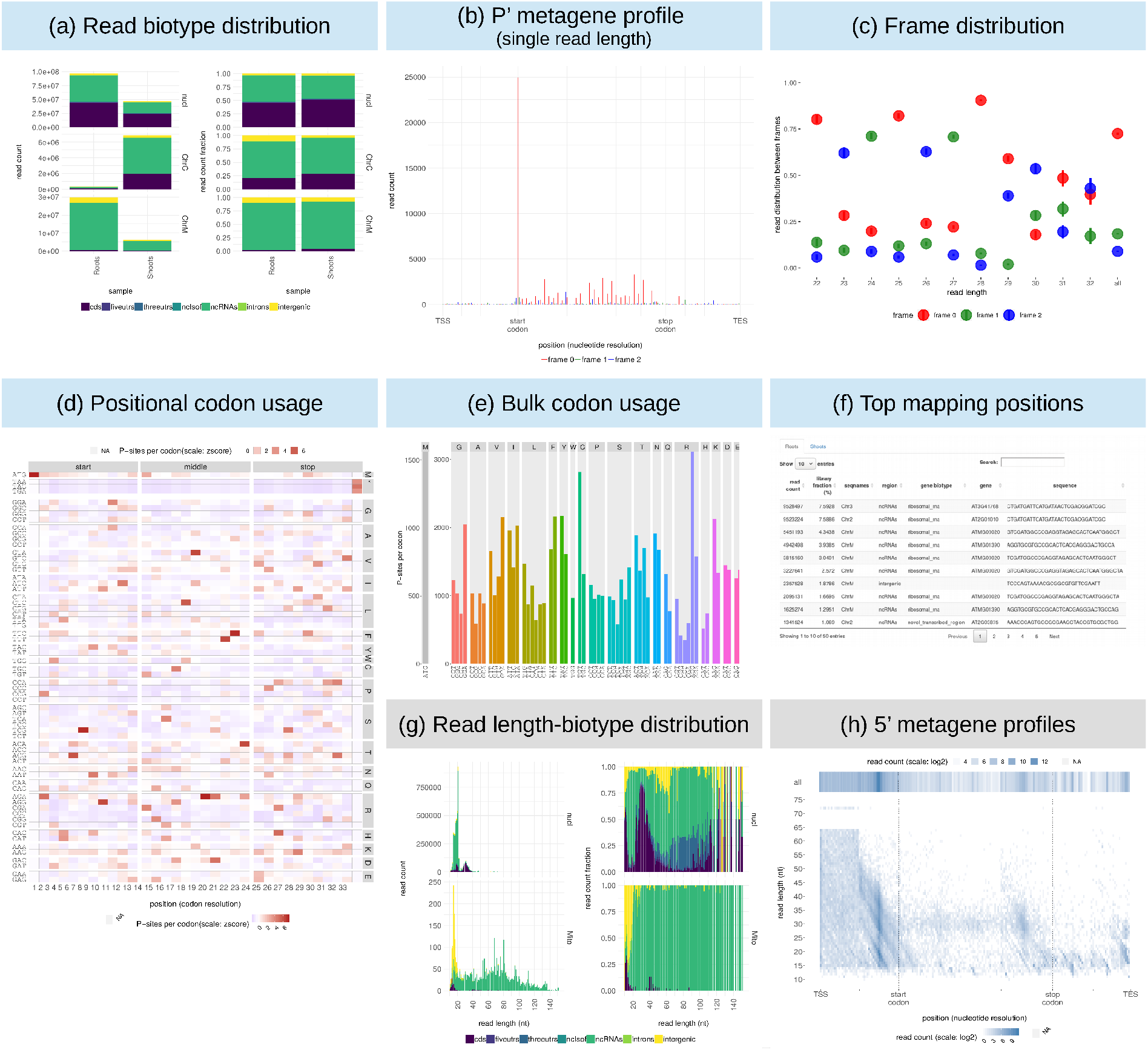
Ribo-seQC analysis for *Arabidopsis thaliana* (blue) and TCP-seq (grey) datasets: (a) biotype-specific read distribution, (b) P-site profile for chloroplast footprints of 29nt length, (c) read length-specific frame distribution, (d) positional analysis of codon usage (P-sites per codon), (e) bulk analysis of codon usage, (f) analysis of top mapping positions, (g) combined read length and biotype-specific distribution, and (h) length-specific 5’ profiles.

The importance of performing sample-specific and footprint length-specific analysis is demonstrated by analyzing a TCP-seq dataset (Archer *et al*., 2016): TCP-seq is able to detect different conformations of scanning and elongating ribosomal complexes, corresponding to distinct footprint lengths. As shown in Fig. 1g, a wider range of footprints is detected in the elongating ribosome sample; as shown by our analysis, many footprint lengths (also shown normalized by their abundance) do not map to CDS regions. Metagene profiles for scanning ribosomes (Fig. 1h) show the enrichment of different footprint lengths around the start codon, enabling the quantification of potential different ribosomal complexes around annotated transcript regions.

Taken together, these results illustrate *Ribo-seQC* as a useful analysis platform for Ribo-seq and related RNA-seq-based protocols. More information about the generated reports, together with a more detailed description of each analysis step and usage guidelines can be found at https://github.com/ohlerlab/RiboseQC.

## Acknowledgements

D. Sydow thanks José Maria Munio Acuna and Kristina Kühn (Humboldt-Universität zu Berlin), Reimo Zoschke and Yang Gao (Max Planck Institute of Molecular Plant Physiology) as well as Felix Willmund and Raphael Trösch (Technische Universität Kaiserslautern) for their feedback on the Ribo-seQC report.

