## Supplemental Table 1 for "Ribo-seQC: comprehensive analysis of cytoplasmic and organellar ribosome profiling data"

### Supplementary information

**Table 1**: Comparison of functionalities offered by selected published tools for Ribo-seq data quality control and visualization.

| | *Ribo-seQc* | *Shoe laces*[1] | *riboviz*[2] | *ribo Waltz*[3] | *Ribo-TISH*[4] | *Ribo Profiling*[5] | *system PipeR*[6] | *RUST*[7] | *ribo SeqR*[8] |
|---|---|---|---|---|---|---|---|---|---|
| Automated standalone report | x | - | - | - | x | - | - | - | - |
| Side-by-side analysis of nuclear & organellar genome | x | - | - | - | - | - | - | - | - |
| Side-by-side analysis of multiple datasets | x | x | - | - | - | - | x | - | - |
| Read length distribution | x | x | x | x | x | x | - | x | x** |
| Read biotype distribution (biotypes covering the whole genome) | x | - | - | x | - | x | x | - | x** |
| Read length and biotype distribution | x | - | - | - | - | - | - | - | - |
| Codon usage (transcriptome-wide) | x | - | - | x | - | x | - | - | - |
| Codon usage (position-specific) | x | - | - | - | - | - | - | - | - |
| 5’ profiles for wide read length range (heatmap) | x | - | - | x | - | - | - | - | - |
| P-site profiles for wide read length range (heatmap) | x | - | - | x | - | - | - | x* | - |
| Automated selection of P-site read lengths | x | x | ? | x | x | x | - | - | - |
| Automated per read length P-site offset calculation | x | x | ? | - | - | - | - | - | - |
| Frame distribution | x | x | - | x | x | - | - | x | x |
| Per transcript 5'/P-site profiles | - | x | x | - | - | x | - | - | x |

* A- instead of P-site profiles
** Shown in publication, but not documented in package.
